# Differential expression and co-expression reveal cell types relevant to rare disease phenotypes

**DOI:** 10.1101/2024.03.12.584594

**Authors:** Sergio Alías-Segura, Florencio Pazos, Monica Chagoyen

**Affiliations:** Computational Systems Biology Group, Centro Nacional de Biotecnología (CNB-CSIC) 28049 Madrid, Spain; Department of Molecular Biology and Biochemistry, Science Faculty, University of Málaga 29071 Málaga, Spain

**Keywords:** scRNA-seq, disease phenotypes, co-expression, differential expression, rare diseases

## Abstract

Disease phenotypes, serving as valuable descriptors for delineating the spectrum of human pathologies, play a critical role in understanding disease mechanisms. Integration of these phenotypes with single-cell RNA sequencing (scRNA-seq) data facilitates the elucidation of potential associations between phenotypes and specific cell types underlying them, which sheds light on the underlying physiological processes related to these phenotypes. In this study, we utilized scRNA-seq data to infer potential associations between rare disease phenotypes and cell types. Differential expression and co-expression analyses of genes linked to abnormal phenotypes were employed as metrics to identify the involved cell types. Comparative assessments were made against existing phenotype-cell type associations documented in the literature. Our findings underscore the utility of differential expression and co-expression analyses in identifying significant relationships. Moreover, co-expression analysis unveils cell types potentially linked to abnormal phenotypes not extensively characterized in prior studies.

**Key points:**

- Cell types underling rare disease phenotypes remain largely unknown
- Single-cell RNA-seq data from healthy tissues can be analyzed to reveal these cell types
- We employed differential expression and co-expression analysis to identify cell types associated with rare disease phenotypes
- We validated our results with known relations described in the literature

## Introduction

While known causative mutations of genetic diseases typically originate from germline mutations, they clinically manifest in a relatively limited array of organs or tissues. Various molecular mechanisms can account for these tissue-specific manifestations (Hekselman & Yeger-Lotem, 2020). For instance, differential gene expression, indicated by varying expression levels across different tissues (determined through bulk RNA-seq), could elucidate a subset of these tissue-specific disease manifestations (Feiglin et al., 2017). Additionally, tissue-specific protein interactions (Barshir et al., 2014) and involvement in tissue-specific biological processes (Sharon et al., 2022) can also explain several cases.

Phenotypes associated with diseases, commonly referred to as clinical signs, comprise a collection of descriptions of pathological manifestations. Phenotypes are particularly pertinent for the clinical characterization of patients with rare diseases, for which in most cases only a phenotypic profile is available, and not a diagnosed disease (Marwaha et al., 2022). Due to the low prevalence of rare diseases, phenotypes play a crucial role in ensuring accurate diagnosis and patient stratification. From a systemic standpoint, leveraging these phenotypes constitutes a strategy for prioritizing patient variants (Kelly et al., 2022) and for the development of other phenotype-aware network-based approaches (Ranea et al., 2022). Indeed phenotypes are reflected at the molecular network level to the same extent as diseases (Chagoyen & Pazos, 2016).

Measurement of gene expression at the single-cell level (Tang et al., 2009) can now be used to explain disease manifestations in terms of cell types, rather than tissues. These data are increasingly used to study mechanisms at the cell level for both common and rare diseases, e.g. (Auerbach et al., 2021; Dobie et al., 2022; Hu et al., 2020; Khan et al., 2020; Öz et al., 2022). They are also used to develop systemic approaches to identify cell types relevant to disease states. For example, Jagadeesh et al. (2022) and Jia et al. (2022) integrated scRNA-seq data with GWAS data to infer the cellular types in which variants cause common disease; and Hekselman et al. (2024) exploited specific gene expression in a cell type to infer the cell type associated with several Mendelian diseases

In this study, our aim is to evaluate the extent to which gene differential expression and co-expression, inferred from single-cell data of healthy tissues, can predict the cell type involved in a particular phenotype across a set of (often unrelated) genetic diseases. This is particularly relevant for rare disease investigation as scRNA-seq cannot be widely used due to cost constraints, technical limitations, and difficulties in sample acquisition. Our hypothesis posits that phenotype genes exhibit higher levels of expression in the cell type involved compared to other cell types within the tissue, or alternatively they are highly co-expressed. We compare our findings with known phenotype-cell associations documented in the literature (Pazos et al., 2022). Differential expression analysis revealed 13% of the known associations, while co-expression analysis elucidated 42% of these associations, potentially uncovering others not currently documented. Furthermore, we assessed the predictive value of both approaches, revealing distinct trade-offs in terms of sensitivity and selectivity, or TPR (true positive rate)/TNR (true negative rate) balance, between the two methods.

## Results

### Methods overview

In this work, we used single-cell RNA-seq data from healthy tissues to compute differential expression and co-expression of genes for each cluster of cells within a tissue. For each tissue we selected relevant phenotypes for analysis based on their anatomical locations according to the Human Phenotype Ontology (HPO) (Robinson et al., 2008). For each cell cluster within a tissue, we performed Kolmogórov-Smirnov statistical tests and corrected for multiple testing to ascertain the extent to which differential expression and co-expression of genes associated with a tissue phenotype are significantly different to the background distribution. Finally we compared our results with a set of phenotype-cell type associations compiled from the literature (see Methods and Figure 1 for overview of the analysis).

**Figure 1.**
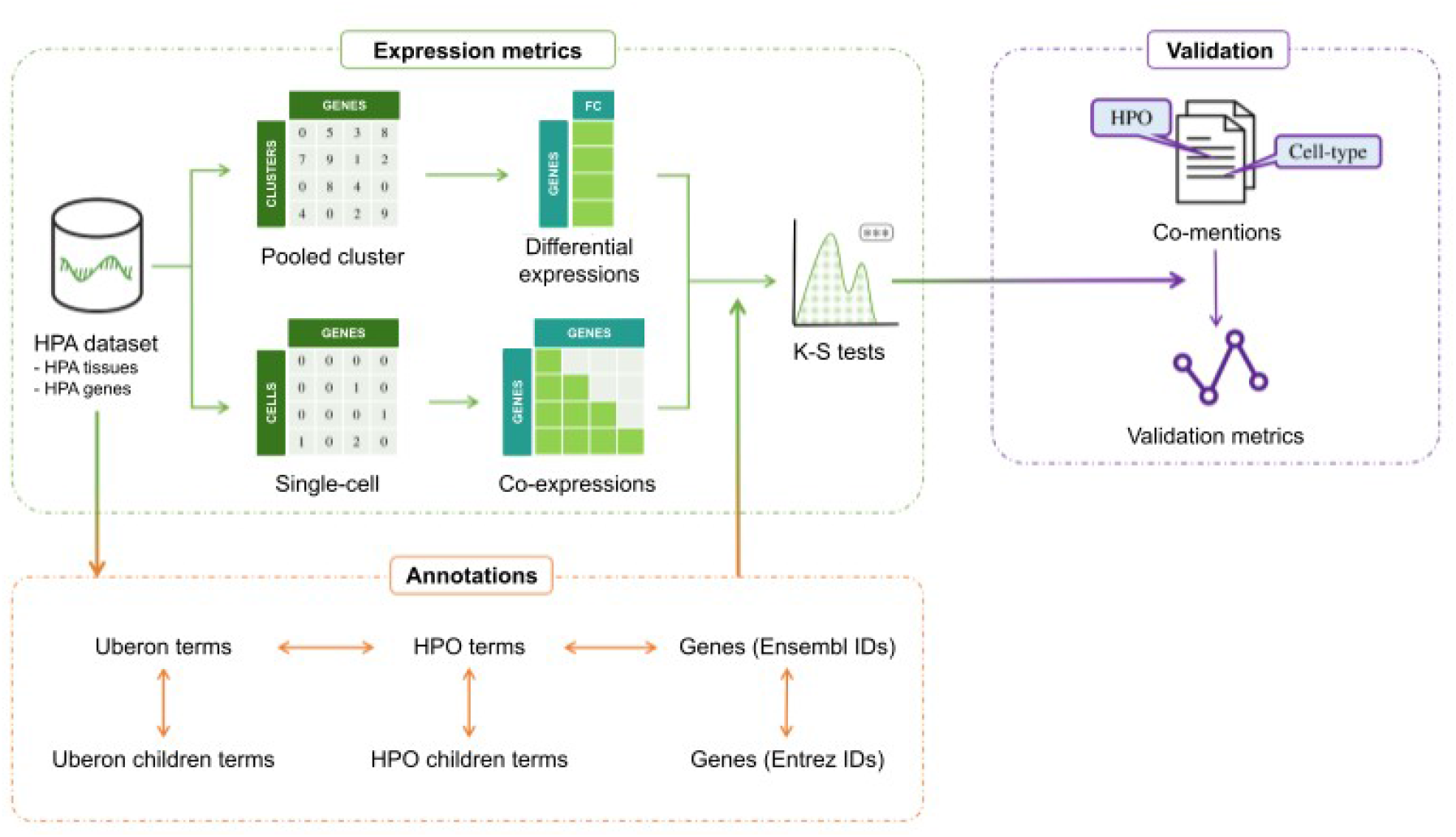
Overview of the analysis. We used single-cell RNA-seq data to compute differential expression and co-expression of genes for each cell cluster within a tissue. We selected relevant phenotypes for analysis based on their anatomical locations according to the HPO ontology. For each cell cluster within a tissue, we performed Kolmogórov-Smirnov statistical tests. We compared our results with a set of phenotype-cell type associations compiled from the literature.

### Cell-type analysis

We analyzed single cell data from 22 tissues compiled from different studies (Bhat-Nakshatri et al., 2021; Chen et al., 2018; De Micheli et al., 2020; Guo et al., 2018; Hawrylycz et al., 2012; He et al., 2020; Henry et al., 2018; Hildreth et al., 2021; Liao et al., 2020; Lukassen et al., 2020; MacParland et al., 2018; Man et al., 2020; Menon et al., 2019; Parikh et al., 2019; Qadir et al., 2020; Solé-Boldo et al., 2020; Vieira Braga et al., 2019; L. Wang et al., 2020; W. Wang et al., 2020; Y. Wang et al., 2020) by the Human Protein Atlas (HPA) (Karlsson et al., 2021), which are annotated with 78 distinct cell-types. After mapping anatomical terms and corresponding HPO terms (see Methods), we finally analyzed 482 non-redundant phenotypes (corresponding to the most specific terms in the HPO hierarchy within those with at least 20 genes) in the context of 444 single cell clusters. The number of phenotypes per tissue finally analyzed is variable (Figure 2). A total of 3,338 phenotype-cell type pairs were analyzed (Table S1).

**Figure 2.**
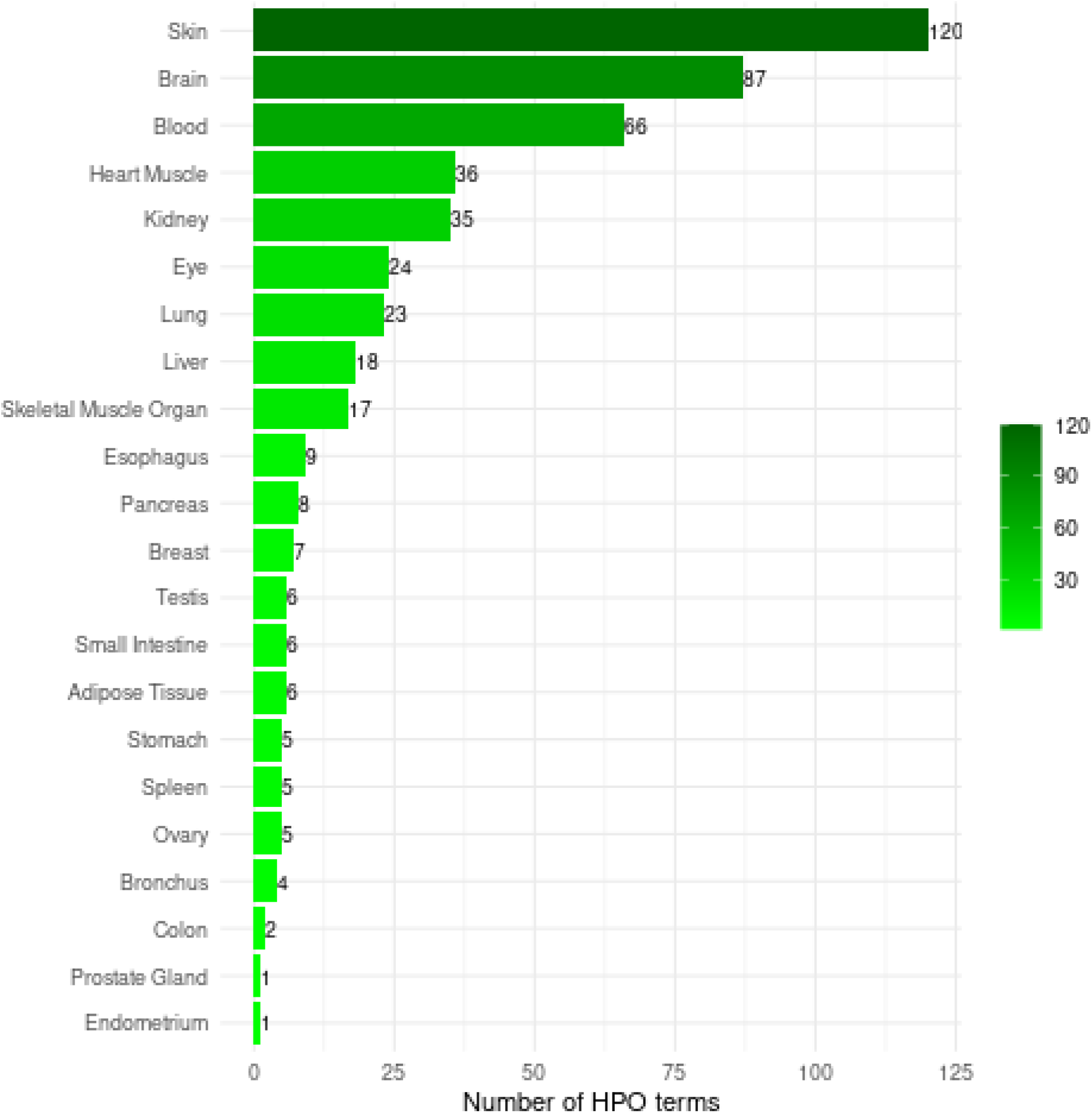
Number of HPO terms analyzed per tissue.

Based on differential expression, we found significant associations (FDR<0.001) for 202 phenotype-cell type pairs. For example, granulosa cells (GCs) from ovary tissue were significantly associated with premature ovarian insufficiency (POI) (Figure 3a). This result is relevant as granulosa cells surrounding oocytes play a pivotal role in folliculogenesis, emerging as an important etiological factor in POI (Liu et al., 2023).

**Figure 3.**
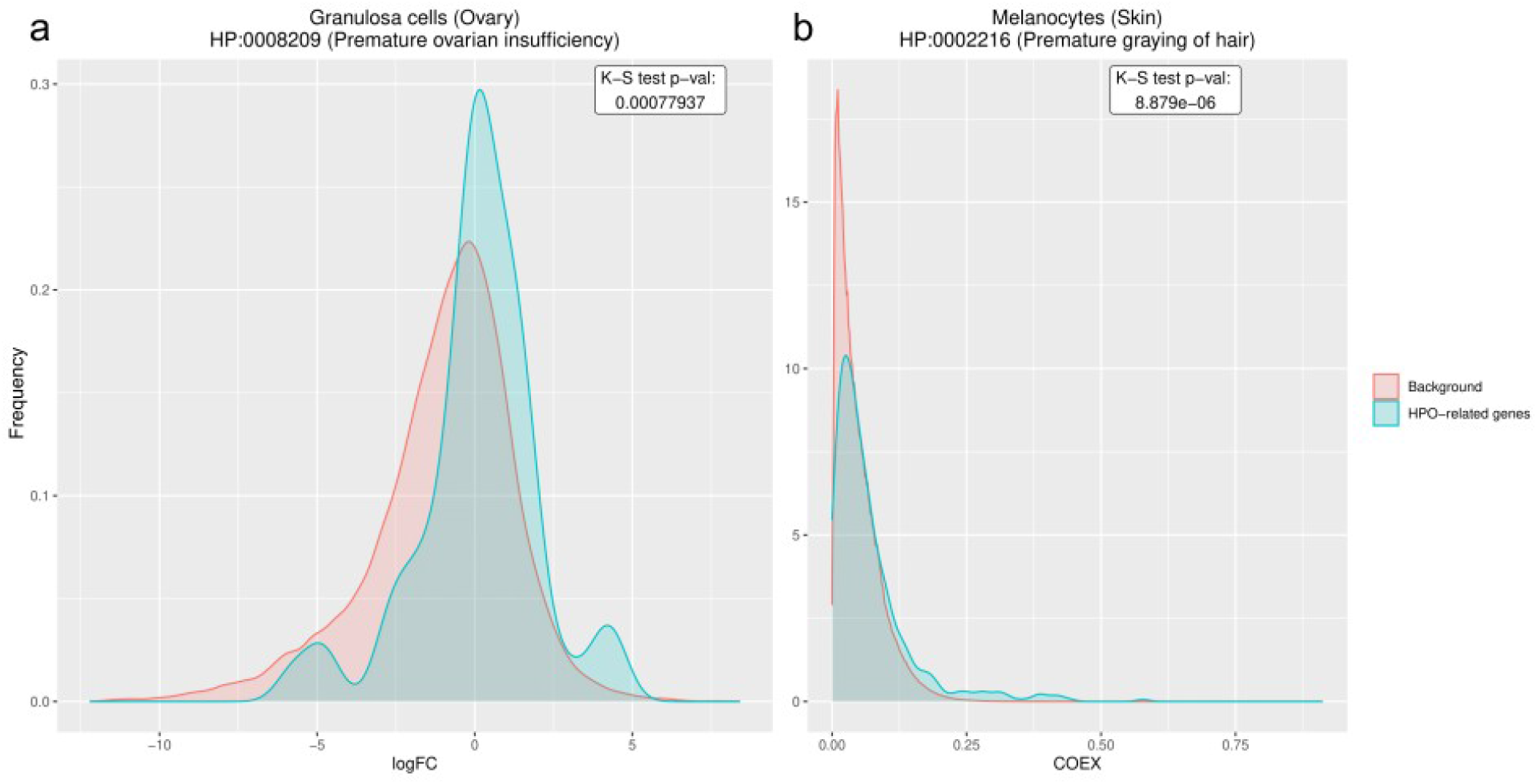
Examples of significant phenotype - cell type associations based on differential expression and co-expression. a. Distributions of FC values of genes associated and not associated with ‘premature ovarian insufficiency’ phenotype in granulosa cells from ovary. The phenotype - cell type association is significant (genes are differentially expressed, P 0.00078). b. Distributions of co-expression values of genes associated and not associated with ‘premature graying of hair’ phenotype in melanocytes from skin. The phenotype-cell type association is significant (there is a higher gene co-expression, P 8.879e-6).

Based on co-expression, we found significant associations (FDR<0.001) for 1,047 phenotype-cell type pairs. As an example, melanocytes from skin tissue were significantly associated with premature graying of hair (Figure 3b). Melanocytes are responsible for producing melanin, the pigment that gives hair its color. Both oxidative stress and genetic factors contribute to the premature graying of hair by impairing melanocyte function (Shi et al., 2014).

We next compared the results obtained by differential expression and coexpression. Spearman correlation of -log10(FDR) values is 0.50 (adjusted R-squared 0.21, P 1.1e-117) (Figure S1). A large number of the phenotypes analyzed (404 of 491, 82.3%) have no significant association by differential expression analysis. In contrast, we found at least one significant association for a higher number of phenotypes (317 of 491, 64.6%) by coexpression analysis. 177 phenotype-cell type pairs were predicted both by differential expression and co-expression, (∼88% of those predicted by differential expression are also predicted by co-expression). For example, proximal tubular cells are significantly associated with proximal tubulopathy (the dysfunction of the proximal tubule, the portion of the duct system of the nephron of the kidney which leads from Bowman’s capsule to the loop of Henle, according to HPO definition), by both differential expression and coexpression.

### Literature validation

To further validate our results and to evaluate the potential predictive value of our approach, we compiled a set of phenotype - cell type associations from the literature using significant co-mentions, following a previously described methodology (CoMent) (Pazos et al., 2022). A total of 541 phenotype - cell type significative associations were considered as positive cases (CoMent p-val<0.001), while those not supported by the literature as negative.

Globally, differential expression can explain 12.75% of the known phenotype - cell type associations (TPR or sensitivity), with a specificity (TNR) of 95.24%. Co-expression achieved a higher sensitivity, explaining 41.59% of the associations, at the expense of a lower specificity (70.61%). (See Table 1 and Table S2 for prediction results at the tissue level).

**Table 1:** Global prediction results.

|  | TP | TN | FP | FN | TPR | TNR | PPV | F1 |
| --- | --- | --- | --- | --- | --- | --- | --- | --- |
| Differential expression | 69 | 2664 | 133 | 472 | 0.1275 | 0.9524 | 0.3416 | 0.1857 |
| Co-expression | 225 | 1975 | 822 | 316 | 0.4159 | 0.7061 | 0.2149 | 0.2834 |

Some of the associations revealed in our analysis might be relevant even if we found no significant co-mentions in the literature. For example, Langerhans cells from skin are associated with recurrent bacterial skin infections by both differential expression and co-expression, and also in some studies (e.g. Rajesh et al., 2019). Basal keratinocytes are associated with skin erosion by differential expression. These cells they are known to possess properties of stem cells and are essential in maintaining the integrity of skin and damage recovery (Yin et al., 2023). And respiratory ciliated cells from bronchus are associated with recurrent bronchitis according to co-expression analysis. It is known that recurrent bronchitis is associated with loss of ciliated cells in children (Gaillard et al., 1994).

## Discussion

In this study, we conducted an exploratory analysis of differential expression and gene co-expression in single cells as an initial step toward investigating the relationships between cell types and rare disease phenotypes. We examined 482 non-redundant phenotypes associated with 22 tissues within the context of 444 single-cell clusters, representing 78 distinct cell types.

Our aim was to identify the specific cell types within a tissue where a phenotype primarily manifests. We deliberately excluded rare disease phenotypes resulting from complex interactions among different tissues. For example, abnormalities in body height (such as short or tall stature) can be attributed to disruptions in cellular functions within both the skeletal and endocrine systems. Therefore, phenotypes not directly associated with a particular anatomical location (as defined in the HPO ontology) were not included in our analysis. With this approach, only nine phenotypes were analyzed in more than one tissue (Table S3), for instance, hepatosplenomegaly, which is related to both the liver and spleen.

Even if manifested within a single tissue, some phenotypes could be caused independently or in association by several cell types within that tissue. According to our validation set 27% of the analyzed phenotypes have more than one cell type discussed in the literature. In some cases, our approach captured the involvement of multiple cell types in a phenotype (Figure S2). For instance, dilated cardiomyopathy is significantly associated with both cardiomyocytes and fibroblasts in the literature (e.g. Tsuru et al., 2023) and both cell types were significantly associated based on both differential expression and co-expression analyses. However, in other cases, partial involvement of a subset of genes in each cell type might not yield significant results for each cell type independently.

Many relationships between phenotypes and cell types remain unknown. According to CoMent, 57% of phenotypes analyzed have no significant cell type association. The percentage is exceptionally high in some tissues, like the brain (69 of 87 phenotypes, nearly 80%, have no significant association according to literature co-mentions). Therefore, our validation set, compiled from the scientific literature, contains a small number of positive cases, implying that the precision (PPV) of our approach is likely underestimated. Predictive approaches like the one described in this work based on healthy tissues can aid in the discovery of the cell types associated with rare disease phenotypes.

Finally, not all genes may be equally relevant for a phenotype, as phenotypes may only occasionally occur in a disease or may be secondary to the cause of the disease. However, in this study, we considered all genes equal, as phenotype frequency is only available for a small number of diseases.

Still with limitations, our results suggest that differential expression and co-expression inferred from single cell data could be used to explore potential cell types involved in the phenotypic manifestations of rare diseases. This knowledge may help in the molecular understanding of rare disease mechanisms, and in the development of improved diagnostic tools and new treatments/therapies.

## Materials and methods

### Single-cell dataset

We obtained single-cell transcript counts (scRNAseq) from 26 healthy tissues from the Human Phenotype Atlas (HPA) Version 21 https://v21.proteinatlas.org/ with their corresponding cluster and cell type annotations (Karlsson et al., 2021). We obtained as well from HPA aggregated transcripts per million (TPM) for each cluster, representing the whole pool of single cells assigned to that cluster.

We decided to exclude some tissues from the analysis due to various reasons. Bone marrow was discarded due to not having HPO terms associated with its Uberon terms. Rectum, lymph node and placenta were excluded due to not having significant literature co-mentions for any of their HPO terms.

### Ontologies

We retrieved disease phenotypes (clinical signs) and their corresponding gene annotations from the Human Phenotype Ontology (HPO) release 2022-02-14 (https://hpo.jax.org) (Robinson et al., 2008). We gathered anatomical entities such as organs, tissues and body parts from the Uber-anatomy ontology (Uberon) release 2022-02-21 (https://www.ebi.ac.uk/ols4/ontologies/uberon) (Mungall et al., 2012). In order to map tissues to phenotypes we proceeded as follows: we manually assigned an Uberon term to every tissue present in the HPA dataset (Table S4), and we retrieved all their descendant terms (more specific terms that also depict the tissue) from the Uberon .obo file. Then, for each Uberon term we obtained the corresponding mappings to HPO from the HPO .owl file. For each of these HPO terms we obtained their descendant terms from the HPO .obo file. Finally, we annotated those HPO terms with their associated genes from the HPO annotation file phenotype_to_genes.txt. We kept only sufficiently informative (with at least 20 associated genes) and sufficiently specific (no descendant terms with more than 20 associated genes) HPO terms for further analysis. In order to annotate HPO terms with their associated genes, we performed Ensembl ID-Entrez ID conversion using BioMart (Kinsella et al., 2011).

### Computing differential expression and co-expression values

We calculated two metrics for each phenotype in each cell cluster (per tissue) based on: differential gene expression and co-expression.

### Differential expression

The HPA dataset aggregated by cluster was used to calculate the differential expression. This dataset contains, for each gene and cluster, a transcripts per million (TPM) value. We computed the fold change (logFC), according to the following formula:

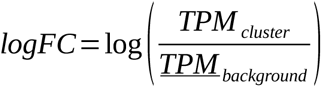

Where TPM_cluster_ is the TPM value for the gene of interest in the cluster of interest, and *<u>TFM</u>*_*background*_is the average TPM value for the gene of interest in the rest of the clusters of the dataset.

### Co-expression of gene pairs

The HPA scRNA-seq dataset was used to calculate gene co-expressions. For each cell cluster in a tissue, we calculated all gene pair co-expressions with COTAN (COexpression Tables ANalysis) (Galfrè et al., 2021). In a pre-processing step, we first removed cell outliers and performed some data quality comprobations adapting the vignette provided by COTAN authors (see Availability for details). To generate co-expression matrices, for each pair of genes we built a 2×2 contingency table containing the number of cells in each possible condition (expressing both genes, only the first, only the second or neither). With this table, we computed the GPA co-expression index (COEX) that measures the deviation of co-expression from the expected proportion under the independence assumption (ranging from -1 to 1).

## Distribution comparisons

For each cell cluster and phenotype pair within a tissue, we employed the Kolmogorov-Smirnov (K-S) test to determine whether genes associated with a phenotype have significantly higher differential expression with respect to the rest of genes or extreme co-expression within them. We finally corrected these significance values (p-values) for multiple testing using the Benjamini-Hochberg False Discovery Rate (FDR) method.

Note that a tissue can have several cell clusters of the same cell type (e.g., tissue liver has 5 clusters annotated as hepatocytes). The final score for a cell type is the minimum FDR of all cell clusters annotated with that cell type. We considered cell types associated with a phenotype if their minimum FDR < 0.001.

## Results validation

We assessed our results with a validation set derived from the literature. For that we built a corpus of literature co-mentions between the phenotypes and cell-types analized using CoMent (Pazos et al., 2022). We considered as known relations those with a CoMent p-value<0.001.

For each of the two metrics studied (logFC, co-expression) we generated a confusion matrix per tissue. Using the confusion matrices, we calculated a set of metrics: true positive rate (TPR), true negative rate (TNR), positive predictive value (PPV), and F-score (F_1_). We similarly calculated the overall confusion matrix and corresponding metrics.

## Availability

The HPA dataset is available at https://v21.proteinatlas.org/about/download, section 9 - RNA single cell type tissue cluster data, and section 10 - RNA single cell read count data. The HPO ontology release 2022-02-14, as well as the annotation file, are available in https://github.com/obophenotype/human-phenotype-ontology/releases/v2022-02-14.

Likewise, the Uberon anatomy ontology release 2022-02-21 is available at https://github.com/obophenotype/uberon/releases/v2022-02-21.

COTAN version 0.99.10 is available from BioConductor version 3.15 (https://bioconductor.org/news/bioc_3_15_release/), that works with R version 4.2. Vignette corresponding to that version is available at the 5d76054 commit from COTAN GitHub repository (https://github.com/seriph78/COTAN/blob/5d760549d0ea7b357dce0ef49d1346619d378217/vignettes/Guided_tutorial.Rmd).

All code generated in this work is available from GitHub at https://github.com/SergioAlias/sc-coex.

## Author’s contributions

S.A.S.: Methodology, software, formal analysis, investigation, writing-original draft, review & editing; F.P.: Data curation, software, validation, writing-review & editing; M.C.: Conceptualization, methodology, formal analysis, investigation, writing-original draft, review & editing.

## Acknowledgements

This work was supported by the Spanish Ministry of Economy and Competitiveness with the European Regional Development Fund [grant numbers PID2019-108096RB-C22 and PID2022-140047OB-C2] to M.C. and F.P.

## AI disclosure statement

During the preparation of this work the authors used ChatGPT 3.5. for the purpose of correcting spelling, grammar and style. After using this tool/service, the authors reviewed and edited the content as needed and take full responsibility for the content of the publication.

## Supplement

**Table S1**. Results for all phenotype-cell type pairs analyzed.

**Table S2**. Prediction results, for differential expression and co-expression, at the tissue level.

**Table S3**. HPO terms analyzed in more than one tissue.

**Table S4**. Manually assigned Uberon term for each tissue.

**Figure S1.**
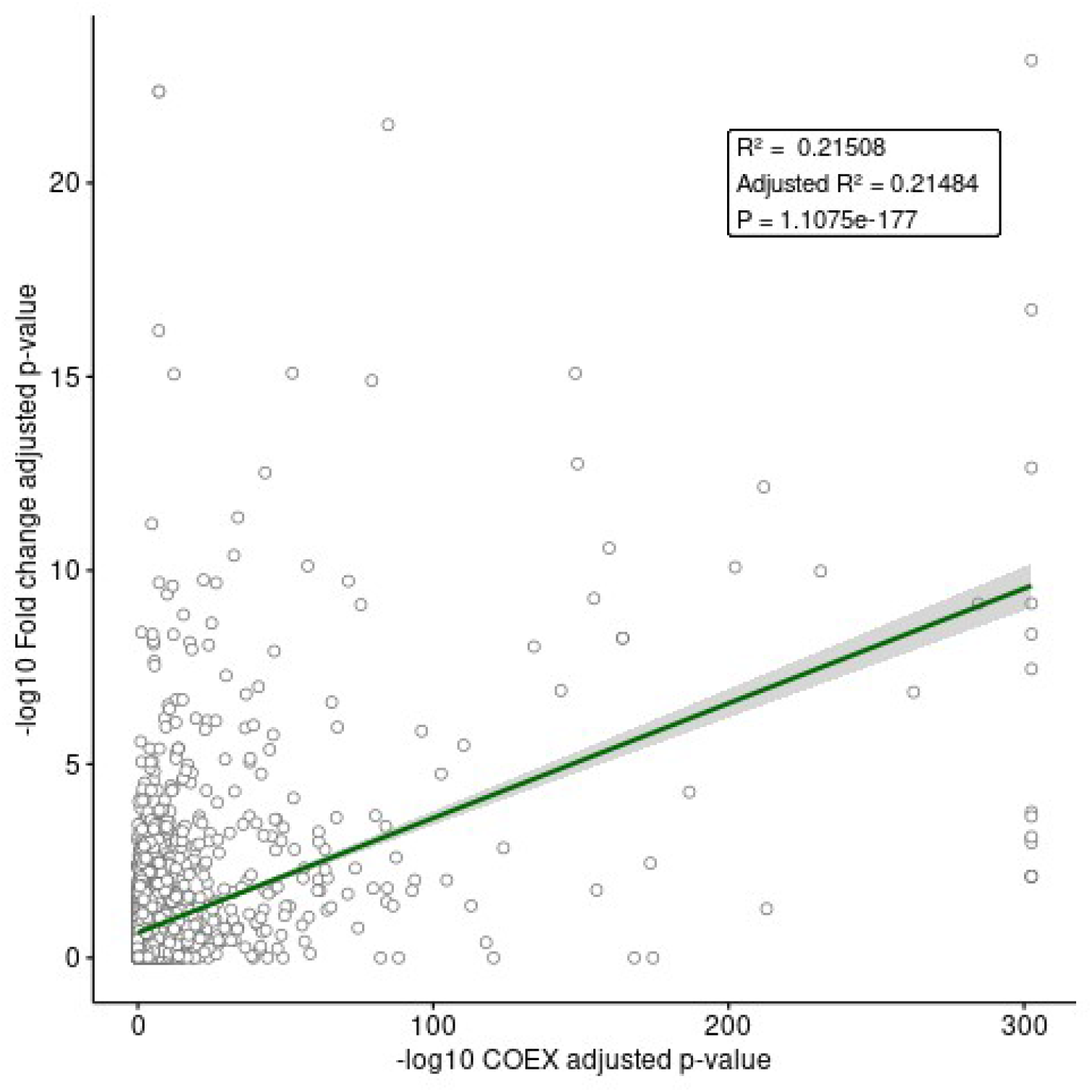
Comparison of differential expression (y-axis) and co-expression (x-axis) results: log-transformed adjusted p-values (FDR) for each phenotype-cell type pair analyzed.

**Figure S2.**
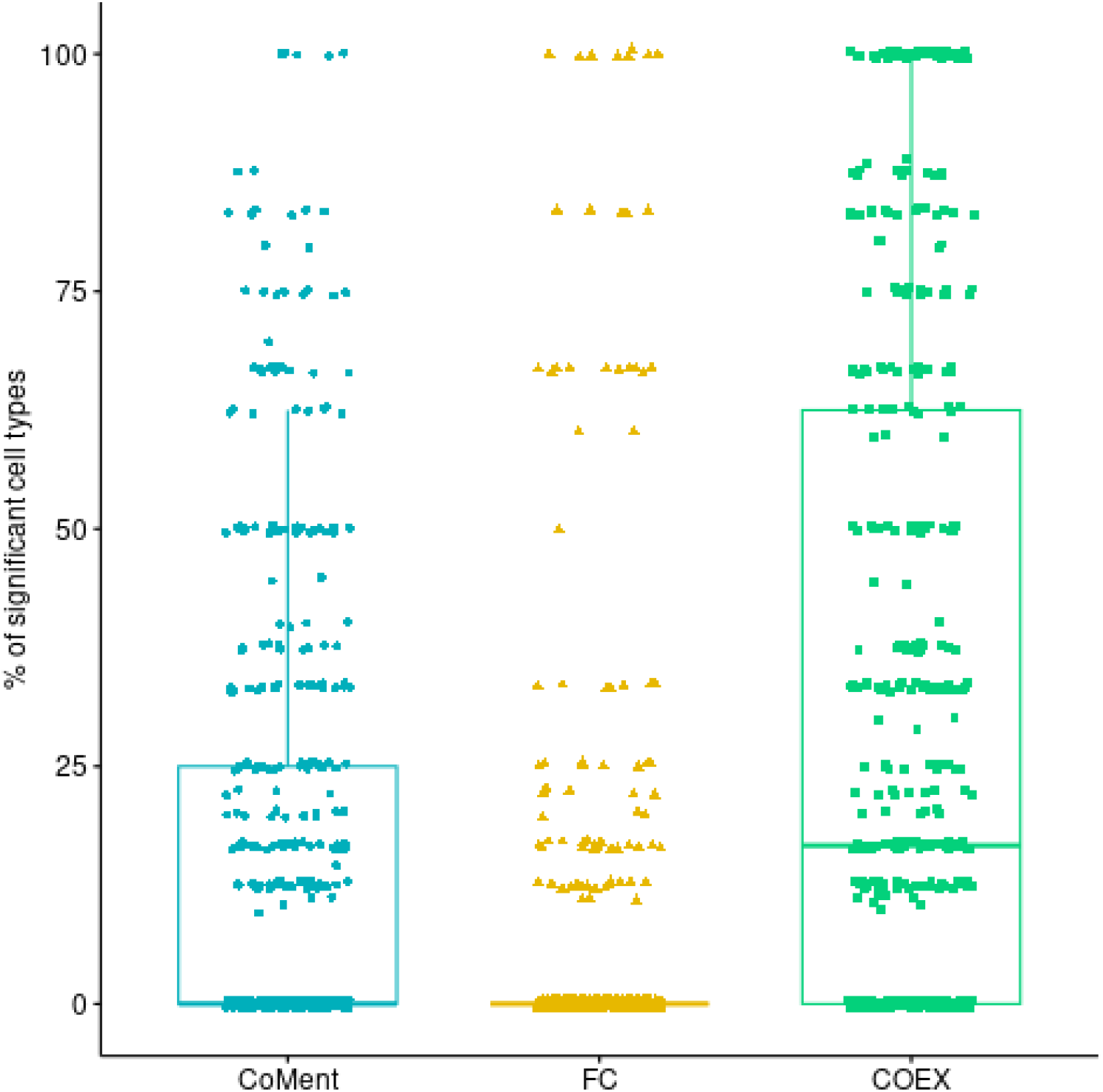
Percentage of significant cell types per phenotype - tissue pair, from literature (CoMent), fold change (FC) and co-expression (COEX).

